# Temporal changes in metabolism late in seed development affect biomass composition in soybean

**DOI:** 10.1101/2020.10.15.341339

**Authors:** Shrikaar Kambhampati, Jose A. Aznar-Moreno, Sally R. Bailey, Jennifer J. Arp, Kevin L. Chu, Kristin D. Bilyeu, Timothy P. Durrett, Doug K Allen

## Abstract

The inverse correlation between protein and oil production in soybeans is well-documented; however, it has been based primarily on the composition of mature seeds. Though this is the cumulative result of events over the course of soybean seed development, it does not convey information specific to metabolic fluctuations during developmental growth regimes. Maternal nutrient supply via seed coat exudate measurements and metabolite levels within the cotyledon were assessed across development to identify trends in the accumulation of central carbon and nitrogen metabolic intermediates. Active metabolic operation during late seed development was probed through transient labeling with ^13^C substrates. The results indicated: i) a drop in lipid during seed maturation with a concomitant increase in carbohydrates, ii) a transition from seed filling to maturation phase characterized by quantitatively balanced changes in the carbon use and CO_2_ release, iii) changes in measured carbon and nitrogen resources supplied maternally over development, iv) ^13^C metabolites processed through gluconeogenesis towards sustained carbohydrate accumulation as the maternal nutrient supply diminishes, and v) oligosaccharide biosynthetic metabolism during seed coat senescence at maturation. These results highlight temporal engineering targets for altering final biomass composition to increase the value of soybeans and a path to breaking the inverse seed protein and oil correlation.

**One-sentence summary:** Assessment of temporal changes in metabolism during soybean seed development indicated that lipid turnover during maturation contributes carbon for gluconeogenic production of carbohydrates.

## INTRODUCTION

The composition of a seed including protein, oil, and carbohydrate levels establishes its commercial value. In soybean [*Glycine max* (L.) Merr.], storage protein accounts for 35-40% of seed dry weight, with lipids (i.e., oil) accounting for 18-20%, predominantly as triacylglycerol (TAG) (Adams et al., 1983; Collakova et al., 2013; Li et al., 2015). At a value of approximately $40 billion per year, soybean production is second only to corn in contribution from a crop to the US economy; United States Department of Agriculture (USDA), National Agriculture Statistical Services (NASS). Other biomass components, including carbohydrates, are of less market value, and a subset (i.e., raffinose family oligosaccharides (RFOs)) produced late in development cannot be metabolized for energy by monogastric animals. The RFOs include raffinose and stachyose and are considered anti-nutritional components of livestock feed, therefore detracting from seed value. In soybean, breeding efforts that increased protein content have resulted in lower yields (Mello Filho et al., 2004; Singh et al., 2016; Assefa et al., 2018) and the production of less TAG, indicating a tradeoff between protein and both yield and oil in mature seeds. Breaking the inverse correlations to improve the total seed value without compromising yields are unrealized goals of most breeding and biotechnological efforts.

Central carbon metabolic pathways are responsible for the production of storage reserves including lipids, proteins, and carbohydrates in plants. Though the network of central metabolism is highly conserved across species, there is significant diversity in biomass compositions within plant tissues, indicating that flux through the metabolic pathways can vary extensively. For example, the level of lipids in reproductive organs can range from less than 1% in peas and lentils to greater than 70% in pecans and walnuts and up to 88% in mesocarp tissues such as palm (Dyer et al., 2008; Bates and Browse, 2012; Allen et al., 2015). Other organs such as leaves have low levels of lipids (<5%) in the forms of phospho- and galactolipids for membranes and very little storage lipid in the form of TAG (Lin and Oliver, 2008; Chapman et al., 2012). This variation indicates that steps in the metabolic network are pliable with throughput being context-specific across organs, species, and environments (Allen et al., 2015; Allen, 2016). However, resources available to a developing tissue such as a seed are finite, being constrained by the supply and form of exudates from the seed coat of the maternal plant, usually comprised of sugars (i.e., sucrose, glucose, fructose) and amino acids (glutamine, asparagine, alanine) (Rainbird et al., 1984; Fabre and Planchon, 2000; Schwender and Ohlrogge, 2002; Pipolo et al., 2004; Hernández-Sebastià et al., 2005; Allen et al., 2009). Thus, final seed composition, including oil and protein quantities, is a consequence of the availability of received assimilates and flux through enzymatic steps in metabolic pathways (Allen and Young, 2013; Truong et al., 2013). Understanding the differences in metabolic network flux and operation in tissues and species provides a template to engineer seeds or other organs with value-added compositions.

A quantitative description of temporal changes in metabolic network operation requires experimental methods that can probe stages of seed metabolism precisely and dynamically. Metabolite levels of primary intermediates such as amino acids, sugars, and organic acids decline throughout development while the storage components that include RFOs, lipids, and proteins increase (Fait et al., 2006; Collakova et al., 2013; Li et al., 2015). These levels, however, are routinely reported on a “per gram” basis. As the content of storage components that are considered “inactive/inert pools” in developing seeds increase, the primary metabolites, i.e., “active pools”, are diluted as indicatated by the hypothetical description (Figure 1A). Hence reports of metabolite levels must properly account for dilution due to reserve accumulation when comparing trends over development. Metabolite amounts when compared on a “per seed” basis take into account the increase in storage reserves and may portray more accurately the transient changes in accumulation (Figure 1B vs 1C).

**Figure 1:**
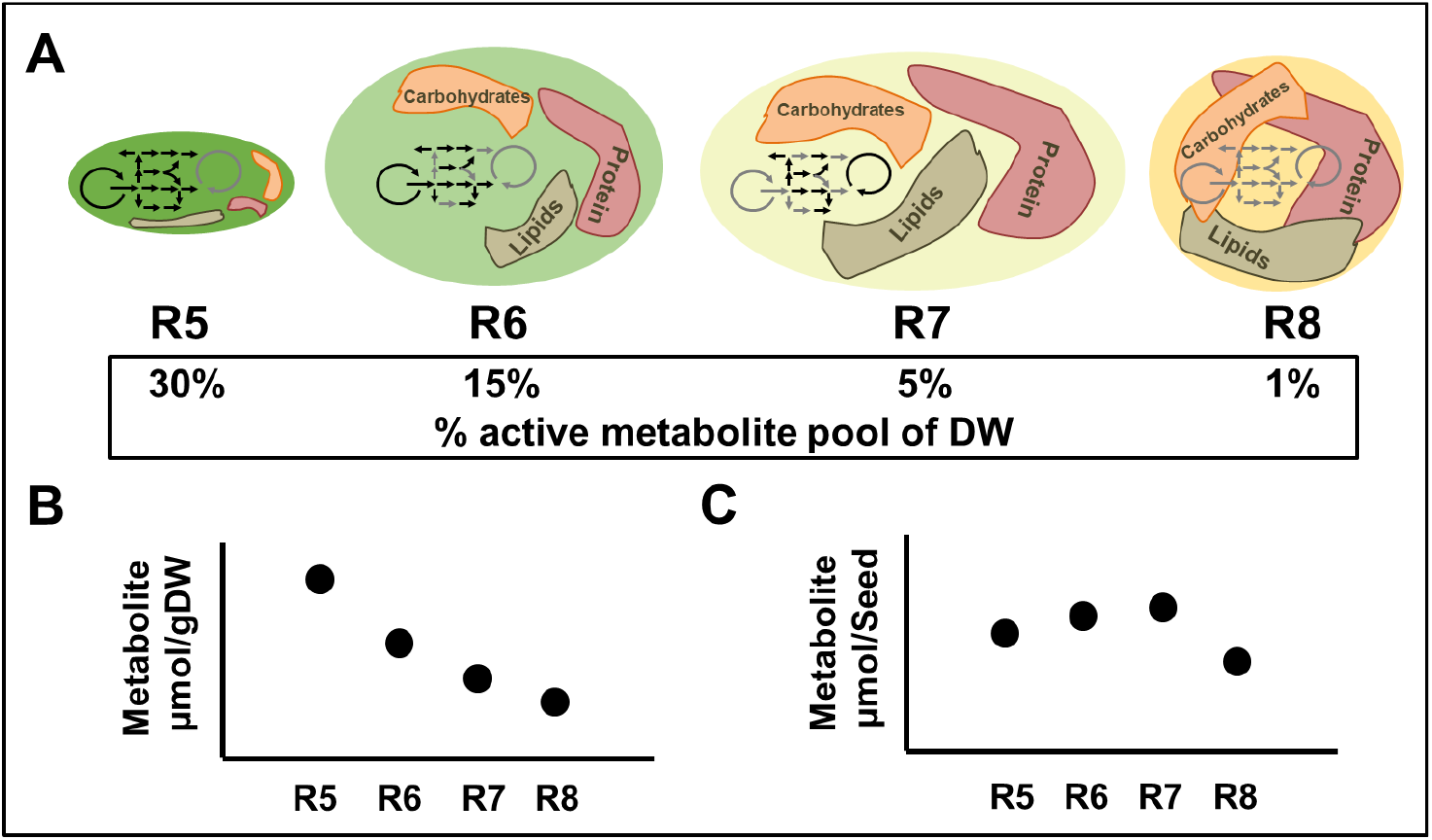
Description of temporal changes in metabolite content over seed development: A. Representation of the increasing accumulation of inactive/inert pools that constitute key storage reserves over the course of development (R5-R8) diluting the active metabolite pool. Decrease in active metabolite content as a percent of biomass (dry weight basis) by developmental stage B. Trend of the active metabolite content as evaluated on a “per gram dry weight” basis C. Trend in the active metabolite content as evaluated on a “per seed dry weight” basis.

One understudied developmental phase of metabolism is seed maturation. The process of desiccation involves more than drying, as indicated by enhanced enzyme activities and gene expression levels (Angelovici et al., 2010), and has important consequences on final reserve composition. However, hypotheses suggested by gene expression and final compositions require validation. Seed maturation represents ~40% of the entire seed developmental progression (Leprince et al., 2016) during which 10-15% of TAGs are turned over (Chia et al., 2005; Baud and Lepiniec, 2009; Baud et al., 2009) and coincidentally, carbohydrates such as RFOs and cell wall polysaccharides (CWPs) continue to accumulate. The metabolic fate of turned over lipid carbon is not clear, but as the supply of exogenous substrates from the maternal plant ceases, sources of carbon are needed to support biosynthetic demands (Baud et al., 2002; Baud and Graham, 2006; Angelovici et al., 2010). Genes involved in fatty acid oxidation and the glyoxylate cycle are expressed at higher levels late in development (Chia et al., 2005; Fait et al., 2006), suggesting that altered tricarboxylic acid (TCA) cycle metabolism, which can vary extensively in seeds (Schwender et al., 2006; Alonso et al., 2007; Allen et al., 2009), might be necessary to meet differing demands (Rolletschek et al., 2003; Rolletschek et al., 2005; Tschiersch et al., 2011) when seed-based photosynthetic contributions decline (Borisjuk et al., 2005; Fait et al., 2006; Angelovici et al., 2010). Whether changes in mitochondrial respiration (Chia et al. 2005) and peroxisomal metabolism (Salon et al., 1988; Raymond et al., 1992; Eastmond et al., 2000; Eastmond and Graham, 2001) could explain repartitioning of carbon for demand late in seed development and support RFO and CWP production (Kuo et al., 1988; Sánchez-Mata et al., 1998; Fait et al., 2006; Collakova et al., 2013; Gawłowska et al., 2017) is unknown.

The result of lipid decreases and production of RFOs during maturation metabolism is a less valuable grain. Experimental results presented here suggest that turned over reserves, including lipids, provide carbon late in seed development necessary to sustain production of RFOs and CWPs. Temporal changes in seed biomass components, the maternal nutrient contribution, and concentrations of central carbon and nitrogen metabolic intermediates were used to study the changing operation of the metabolic network during seed development. Stable isotopes were used to probe cotyledon and seed coat metabolism specific to the maturation phase to describe changes in carbon partitioning late in development. The differences suggest an engineering opportunity to make seeds with value-added composition by paying attention to temporal effects.

## Results

### Changes in soybean seed biomass composition during maturation decrease seed value

Seeds were harvested according to size based on contemporary developmental stage descriptions (Naeve, 2005; Licht, 2014) (Figure 2A), removed from pods and seed coats, and weighed (fresh weight; FW). Cotyledons were dried to determine dry weight (DW) and moisture content (Figure 2B). R5 seeds were comprised of 81.5 ± 2.5% moisture, with seed desiccation events reducing this amount to 53.4 ± 0.5 % in R7, 45.9 ± 1.6 % in R7.5, and 12.9 ± 0.8 % in R8 (maturity). Further loss of moisture continued in mature seed over time to less than 9% dry weight categorized as R8b and R8c. The net CO_2_ release from cotyledons was quantified and indicated a peak in CO_2_ release at R6 followed by a rapid decline (Figure 2C). The measured CO_2_ spike was consistent with differences in storage reserve production, including significant CO_2_ generation when flux from pyruvate to acetyl-CoA enables fatty acid biosynthesis. The lipid production between R6 and R7 over a two week period accounts for 64% of the CO_2_ generated in this time interval based on the accumulation of lipid in seeds (Figure 3).

**Figure 2:**
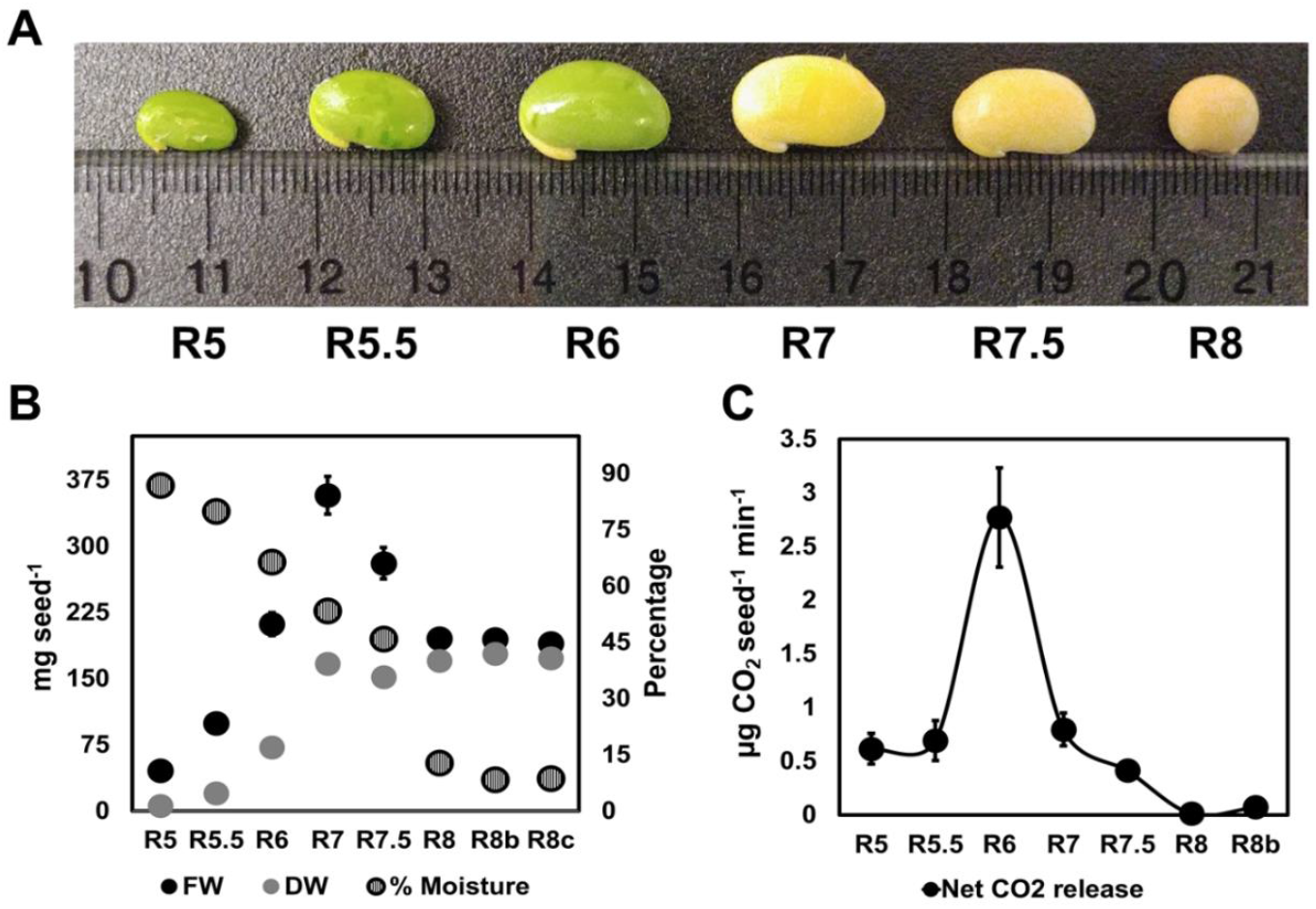
Soybean seed developmental stage descriptions: A. Image of representative cotyledons excised from the seed coat used for analyses from R5 (seed filling stage) – R8 (maturity). B. Fresh weight (FW) and dry weight (DW) measurements of cotyledon pairs represented as mg seed^−1^, with moisture content calculated as loss of water from cotyledons upon drying, are represented in percentages. Error bars represent standard errors of mean, *n* = 6. C. Net CO_2_ released is presented in μg seed^−1^ min^−1^, error bars represent standard errors of mean, *n* = 6 where each of the replicates represents an average of 10 measurements for a single cotyledon to overcome instrument drift.

During maturation of cotyledons there was a small but insignificant drop in total protein accumulatation from 67.6 ± 3.6 mg seed^−1^ at R7 to 59.7 ± 4 mg seed^−1^ at maturity (*p* = 0.11) and a more dramatic significant change in lipid from 40 ± 1.1 mg seed^−1^ at R7 down to 30.4 ± 0.2 mg seed^−1^ at maturity (*p* = 0.004) (Figure 3). Starch production peaked at R6 (4.3 ± 0.2 mg seed^−1^) and declined sharply before leveling off at 0.55 ± 0.1 mg seed^−1^. Sucrose accumulation peaked at 2.2 ± 0.2 mg seed^−1^ during R7 and remained at that level until maturity, while the RFOs raffinose and stachyose accumulated between R6 and R7 to a maximum of 1.6 and 5.9 mg seed^−1^, respectively. Total free amino acids increased modestly throughout development (to 0.6 mg seed^−1^ at R6), and the remaining biomass largely attributed to dietary fiber, including CWP, reached 62.3 ± 7 mg seed^−1^ at R8. All biomass components increased between R5 to R6, indicating that inverse correlations between individual components are not obligatory when sufficient resources were present. Starch turned over between R6 and R7, when significant lipids (67.5%) and RFOs (84.1%) were being synthesized.

**Figure 3:**
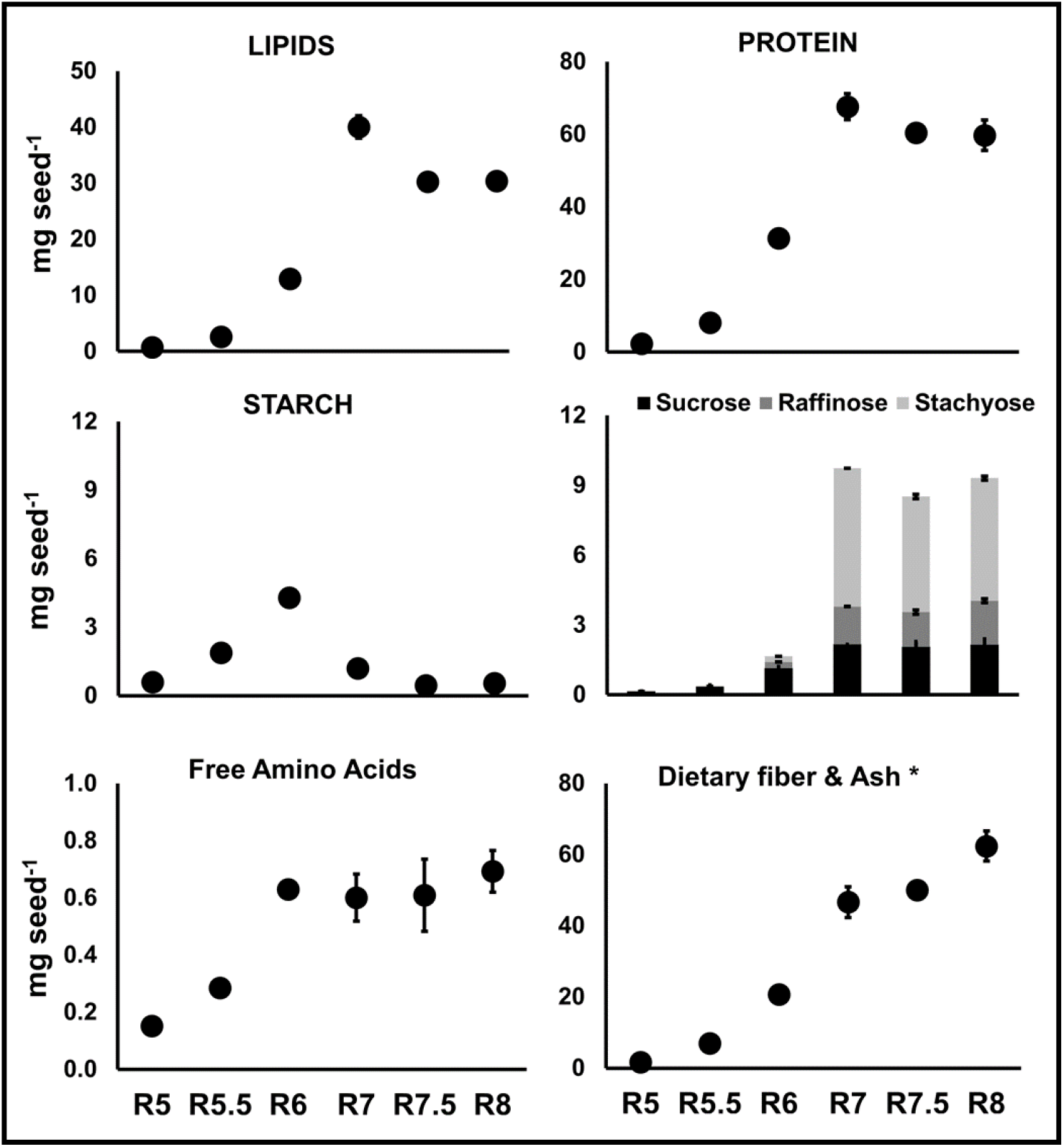
Trends in biomass component accumulation during seed development: Levels of individual biomass components were quantified as described in the ‘Materials and Methods’ section on a mg per seed basis. Values are based on cotyledons and do not include seed coat except for R8. *Dietary fiber and ash were calculated by subtracting all other components from total seed biomass. Error bars represent standard error of mean (*n* = 3).

### Vegetative carbon and nitrogen sources diminish prior to seed maturation in soybeans

Vegetative parts of the plant are the source of sugars and amino acids for developing seeds during much of development (Hsu et al., 1984; Rainbird et al., 1984; Gifford and John, 1985; Egli and Bruening, 2001; Hernández-Sebastià et al., 2005) and impact final seed composition (Allen and Young, 2013); however, the provisions change as seeds mature. Unlike the significant liquid endosperm present in *Brassicaseae*, soybean seed coat exudate is barely detectable at any given stage in development and is present as a shiny wet surface on cotyledons that amounts to a few microliters and diminishes with development. To recover the maximum amount of seed coat exudate for measurements and minimize extraction from within the testa, the surface contents of the interior of the seed coat were briefly extracted with an isotonic solution of 20 mM ammonium acetate, pH 6.5. The exterior surface of the developing cotyledon was also briefly immersed in the same buffer to capture and quantify the major contents of the exudate. An overall decreasing trend of total exudate metabolites (Figure 4) was observed during seed development. The total metabolite levels in the exudate decreased significantly as seeds approached maturation phase (R7 and R7.5), consistent with the trends in reserve accumulation (Figure 3) where no new storage proteins or lipids are made past R7, thus indicating a metabolic transition prior to maturation.

**Figure 4:**
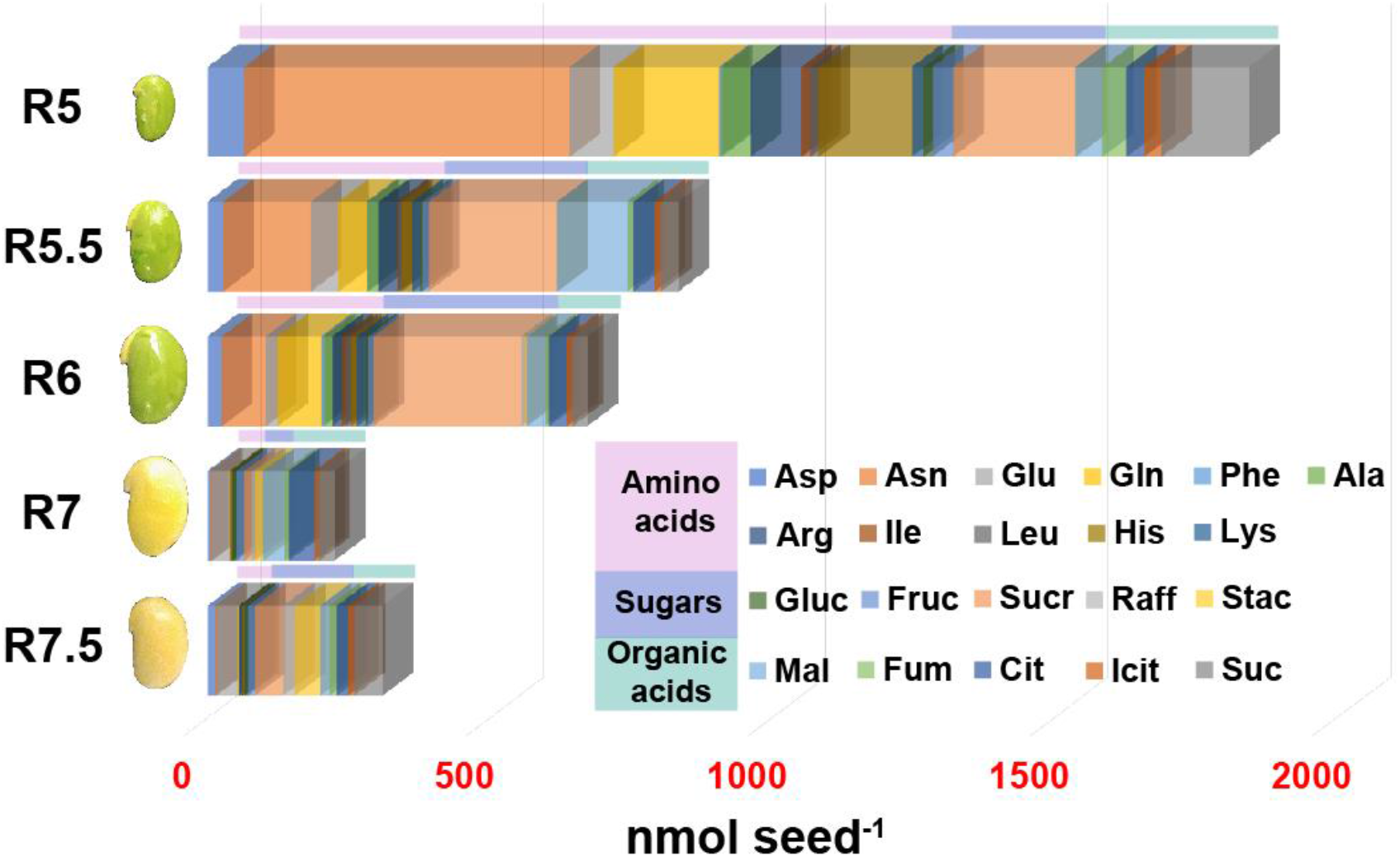
Levels of quantifiable metabolites in the exudate of developing seeds. Three metabolite classes: amino acids (purple), sugars (blue), and organic acids (green) were quantified and presented as nmol per seed amounts. Amino acids represented a major supply in R5 before decreasing significantly (by ~60%) at R5.5. The supply of sugars remained relatively consistent between R5 and R6 and decreased considerably by the time seeds reached the maturation phase (R7). Of the measured metabolites, asparagine (Asn), glutamine (Gln), histidine (His), sucrose (Sucr), malate (Mal) and succinate (Suc) were present in abundance at all stages.

Nine amino acids were measured at detectable levels in the exudate of all development stages. Alanine and lysine were detected in R5, R5.5, and R6. Methionine, threonine, tryptophan, serine, glycine, cysteine, tyrosine, and valine were not detected and must be generated from seed-based metabolism during development (Figure 4, Supplementary Table S1). The nitrogen rich amino acids asparagine, glutamine, arginine, and histidine were among the highest in content during peak protein accumulation stages (R5-R6), consistent with prior studies that indicated glutamine and asparagine are important sources of nitrogen for filling soybeans at least during early stages of development (Rainbird et al., 1984; Hernández-Sebastià et al., 2005). The accumulation of nitrogen-rich amino acids including aspartate, glutamate, and arginine during the last stage of development (from R7.5 to R8) could hint at the importance of nitrogen provision for amino acid biosynthesis during germination.

Sucrose was also a significant carbon source through R6 before decreasing (Figure 4, Supplementary Table S1), possibly due to raffinose and stachyose production from the seed coat at R7 and R7.5 stages. Prior reports that investigated RFOs in young developing seeds (Gomes et al., 2005; Kosina et al., 2009) suggested that the exudate contains precursors to RFO biosynthesis, i.e., sucrose, myo-inositol, chiro-inositol, and pinitol, but did not consider stages of development beyond R6. Data in the current study indicate RFOs are produced and exuded from the seed coat during the maturation phase, i.e., R7 and R7.5 (Figure 4, Supplementary Table S1).

### *In planta* levels of metabolites in cotyledons change throughout seed development

The pathway intermediates that are characteristic of stage-specific metabolism were investigated through pool size quantification (metabolite quantities) in cotyledons. Measurements were first obtained on a “per mg DW” basis and converted to amounts per (dried?) seed to account for the increasing inert pools (lipids, protein, and carbohydrates) over the course of development and enable comparison between different stages (Supplementary Table S2). A *k*-means clustering approach was used to compare changes in trends of metabolite pools over development (Figure 5). All measured amino acids except glutamine clustered into groups 1 and 5, consistent with protein accumulation and a demand for storage protein synthesis plateauing between R7 and R7.5 when storage protein accumulation stopped (Figure 3). The steep increase in amino acid content between R7.5 and R8 (more pronounced for cluster 5) along with the concomitant decrease in storage protein (Figure 3) suggested proteolytic activity during maturation. Cluster 2 consisted of only two metabolites, glucose and fructose, which were elevated at R5 and dropped by R5.5. The variation within this cluster past R5.5 was high, likely due to an increase in glucose content past R6 that was not observed for fructose (Supplementary Table S2). The levels of glucose and fructose in R5 may be a consequence of sucrose breakdown, whereas the increase in glucose at R6 and R7 occured when starch was turning over (Figure 3) supported by the presence of the starch degradation product maltose detected in R6 and R7 (Supplementary Table S2). Maltose, in cluster 3, was similar to the organic acids 2-oxoglutaric acid (2OG), malate, succinate, and fumarate involved in the tricarboxylic acid (TCA) cycle and the sugar phosphates 6-phosphogluconate (6PG), ribose 5-phosphate (R5P), and sedoheptulose 7-phosphate (S7P) involved in the oxidative and reductive steps of pentose phosphate pathway metabolism (PPP). Cluster 3 increased to R6 then declined, similar to measured net CO_2_ release (Figure 2C), suggesting that CO_2_ released in R6 resulted from TCA cycle and OPPP activity along with fatty acid biosynthesis, which is necessary to produce lipids during the same time frame.

**Figure 5:**
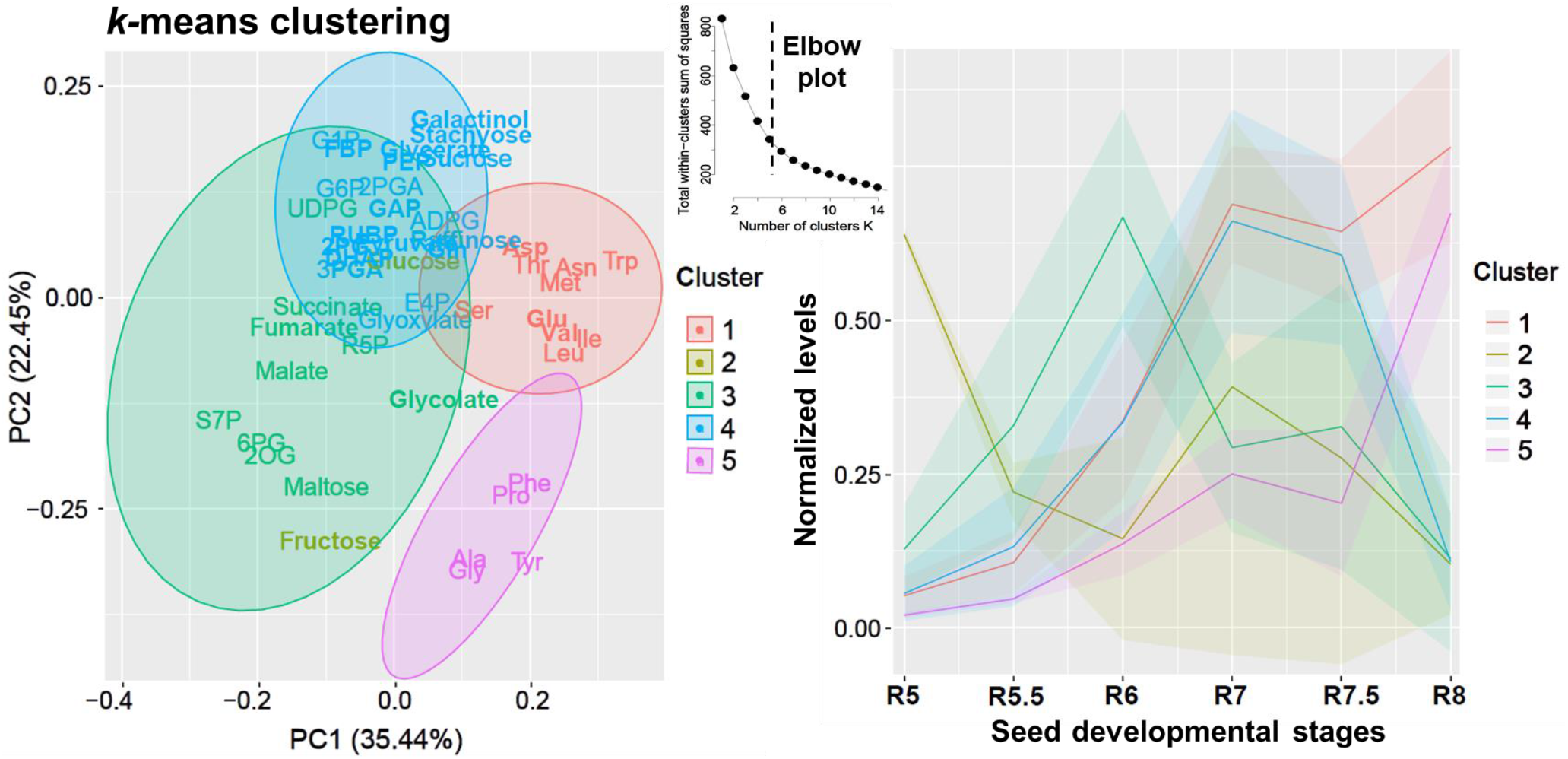
*k*-means clustering using central carbon and nitrogen metabolism intermediates representing trends over seed development stages (R5-R8). A total of 47 metabolites that include central carbon intermediates, organic and amino acids as well as sugars were used for clustering. Metabolite levels were first calculated as nmol seed^−1^ (Supplementary table S2). For the clustering analysis, each metabolite was normalized using its maximum value at any stage (which was given a value of 1) in order to enable comparison of trends over the course of seed development. The optimal number of clusters was determined using the elbow method and was set at *k* = 5, as the within-cluster sum of squared distances reduced past 5 clusters. Metabolites that clustered together were represented on a two-dimentional space using PCA (left panel) and the trends over development for each cluster were presented on the right panel. Abbreviations are defined in Supplementary Table S2.

Metabolites in cluster 3 and cluster 4 (Figure 5) overlapped significantly based on the two-dimentional representation of clusters with principal component analysis (PCA). The common metabolites included those from glycolytic/gluconeogenic pathways and the Calvin-Benson Cycle. Non-overlapping metabolites (sucrose, galactinol, stachyose) were associated with RFO accumulation. The decrease in cluster 4 late in development was consistent with RFO production giving way to CWP biosynthesis (Supplementary Table S2).

### Lipid turnover supports biosynthesis of carbohydrate reserves late in development

Both the supply of resources from the exudate and the metabolic events as indicated from the pool size comparisons and altered storage reserve profiles changed as seeds developed. Pool sizes are suggestive of changes in metabolism but the differences in pool sizes cannot be strictly attributed to altered biosynthetic or turnover rates. Given that protein and lipid levels decrease nearing maturity while CWPs and undesirable RFOs accumulate (Figure 3), isotope tracers were used to probe the movement of carbon into and out of metabolic pools. After validation of the culturing approach (Supplementary Data 1), labeled ^13^C_3_ glycerol was provided to cotyledons for up to 30 minutes to examine metabolism specific to the lipid degradation at the beginning of maturation phase (R7).

R7 seeds incorporated glycerol to produce triose and hexose phosphates over the course of 30 minutes (Figure 6A), consistent with the capacity of operating enzymes gluconeogenically to convert trioses into carbohydrates. Glycerol was chosen in part because entry into metabolism as dihydroxyacetone phosphate (DHAP) would mimic the source of DHAP from the glycerol-3-phosphate backbone that remains after lipolysis late in development. By 30 minutes, labeled carbon originating from ^13^C_3_ glycerol was present at measurable levels in DHAP, GAP, PGA, G6P, G1P, and UDPG (Figure 6A), suggesting that gluconeogenesis may occur late in seed development. Labeling results were further confirmed considering the activity of phosphoenolpyruvate carboxykinase (PEPCK), a key enzyme in gluconeogenic metabolism. PEPCK activity was highest at R7 (Figure 6B), consistent with gluconeogenic activity to supply carbon for carbohydrate metabolism at this stage (Figure 6C).

**Figure 6:**
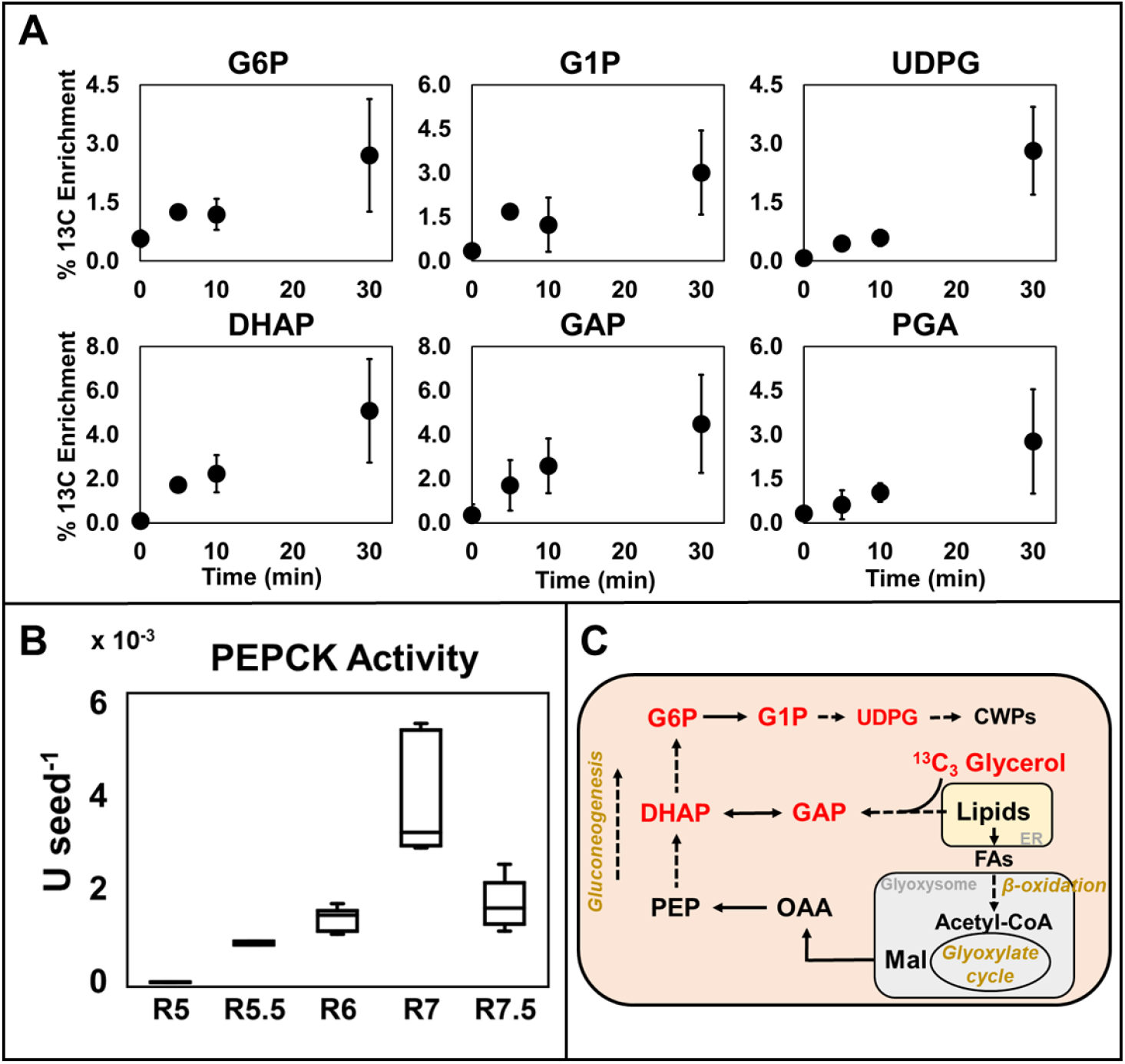
Carbon from turned over lipids is used to make hexose phosphates during R7. A. ^13^C enrichment in intermediates of gluconeogenesis within a 30 min time course pulse labeling experiment using ^13^C_3_ glycerol. B. Phosphoenolpyruvate carboxykinase (PEPCK) activity, as a signature of gluconeogenesis, over the couse of seed development. C. Schematic representation of carbon movement at R7 through central carbon metabolism involved in shuttling carbon from degrading lipids toward carbohydrate metabolism. Intermediates of gluconeogenic and carbohydrate metabolism that were labeled by ^13^C_3_ glycerol are highlighted in red. Abbreviations: G6P, Glucose 6-phosphate; G1P, Glucose 1-phosphate; DHAP, Dihydroxyacetone phosphate; GAP, Glyceraldehyde 3-phosphate; UDPG: Uridine diphosphate glucose; PGA, 3-phospho glyceric acid; PEP, Phosphoenoyl pyruvate; PEPCK, Phosphoenoel pyruvate carboxykinase; OAA, Oxaloacetic acid; Mal, Malic acid; FA, Fatty acids; ER, Endoplasmic reticulum; CWP, Cell wall polysaccharides.

### Isotopic labeling of seed coats indicates production of oligosaccharides that are partitioned to the surface of maturing cotyledons

The presence of RFOs in the exudate was unanticipated and may reflect biosynthetic activities in the seed coat. RFOs in seeds are not required for desication tolerance or germination (Dierking and Bilyeu, 2009; Valentine et al., 2017), and metabolism of the seed coat during development has not been previously described; thus the role of these oligosaccharides remains obscure. During development, the seed coat dry weight is reduced with age, indicating that it may be partially remobilized as the last filial tissue that provides reserves to the seed or could be helpful as an osmotic regulator during germination and the imbibition process. To test the contribution of the seed coat, detached seed coats from the R7 stage were labeled with ^13^C-sucrose, resulting in the production of ^13^C-raffinose (see methods for culture system set up) (Figure 7A, B). The soybean cultivar ‘Jack’ and a near isogenic ultra-low RFO line (Jack *rs2 rs3*) that contained natural variations in the two *raffinose synthase* genes *RS2* and *RS3* were used instead of Williams 82 due to the availability of the normal/ultra-low RFO near isogenic line pair (see methods for details) (Hagely et al., 2020). As shown in Figure 7C, a significant amount of ^13^C incorporation into raffinose within the seed coat was observed in the WT line Jack relative to the ultra-low RFO line (Jack *rs2 rs3*). This result indicated unequivocally that RFOs result in part from seed coat metabolism and may be an important engineering target to favorably alter soybean seed composition.

**Figure 7:**
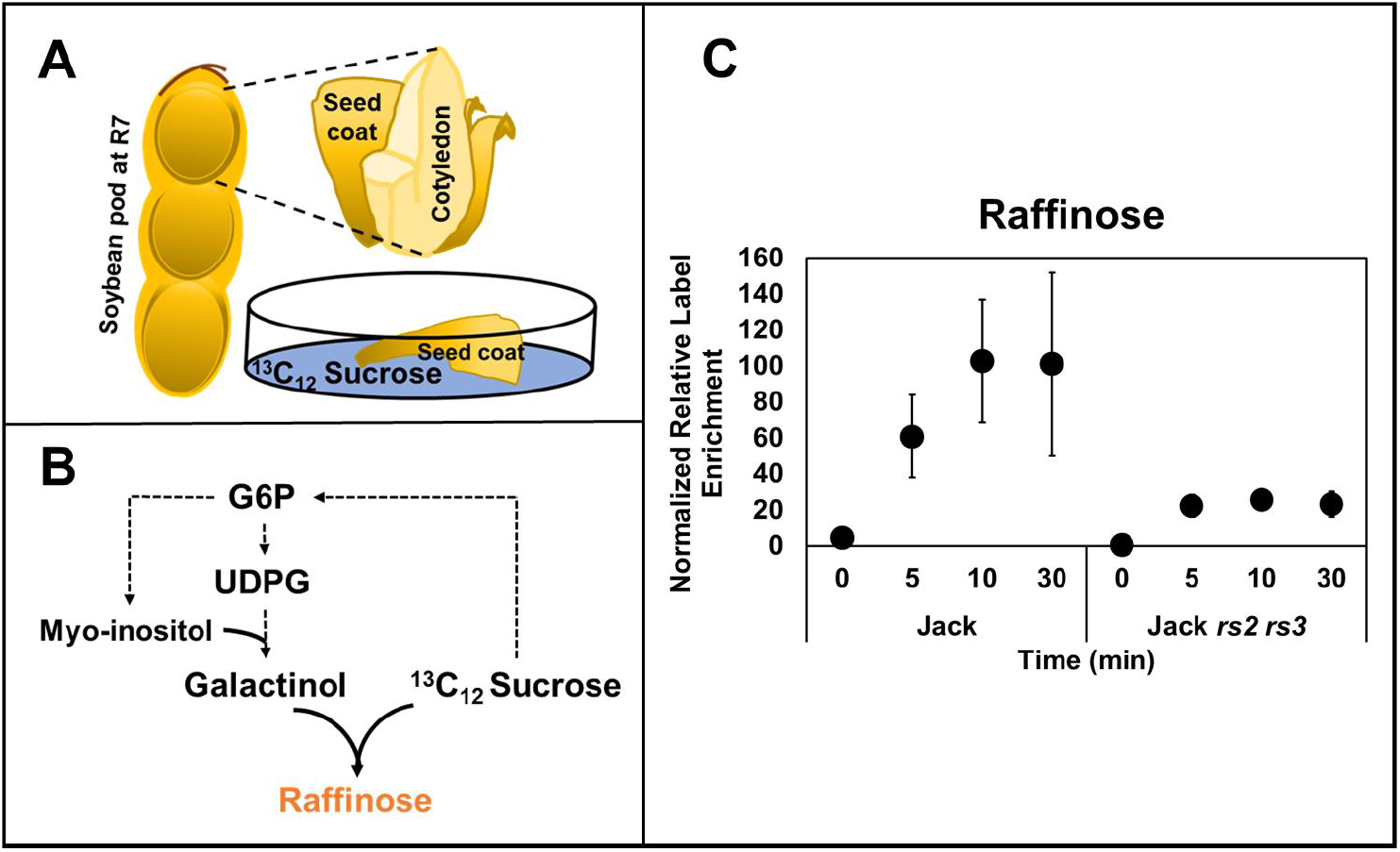
^13^C_12_ Sucrose labeling in seed coats of R7 seeds. A. Depiction of R7 pod with an expanded view of seed coat and cotyledon. Seed coats were excised and cultured with ^13^C_12_ sucrose as a sole source of carbon (see methods for description) over 30 minutes. B. Biochemical route for ^13^C_12_ sucrose incorporation for raffinose biosynthesis. Sucrose is used for the production of glucose 6-phosphate (G6P) followed by myo-inositol. G6P enters carbohydrate metabolism to produce uridine diphosphate glucose (UDPG). UDPG and myo-inositol together produce galactinol which is combined with sucrose to produce raffinose via *raffinose synthase* (*RS*). C. A 30 minute pulse labeling experiment using seed coats of the soybean line ‘Jack’ and a near isogenic ultra-low RFO line, Jack *rs2 rs3* (Hagely et al., 2020) at the initiation of maturation stage (R7) incubated with ^13^C_12_ sucrose indicated significant label enrichment in raffinose. Y-axis represents arbitrary values normalized for pool size comparisons (see Supplementary Table S9) due to significantly different pool sizes of raffinose allowing for direct comparison of label (^13^C) enrichment between the two genotypes. Error bars represent standard error of mean (*n* = 3).

## Discussion

Mature seed composition is the cummulative effect of events throughout development and is a consequence of: a) supply of carbon, nitrogen, and other resources from the maternal plant, and b) seed-based metabolism. In this study we probed the changes in composition over seed development to explain the reduction in oil and protein levels and the accumulation of oligosaccharides late in development. Though inverse protein to oil relationships have been reported regularly (reviewed in Clemente and Cahoon, 2009; Patil et al., 2017), the levels of these two reserves are not at odds during development (Kambhampati et al., 2019). Levels of both lipid and protein in the seed are highest during the initiation of the maturation phase and coincidentally decline at the time when CWPs are the only biomass component being accumulated (Figure 3) when little exogenous carbon is available (Figure 4) to support biosynthetic demands. Thus, the pull for carbon is not exclusive to oil and protein and their turnover is likely a source for production of other reserves.

### Seed-based carbon use efficiency indicates the redistribution of reserves to support metabolism late in development

Carbon conversion efficiency or carbon use efficiency (CUE) has been described in seeds with flux analyses to account for the production of CO_2_ relative to substrates taken up (Schwender et al., 2004; Alonso et al., 2007; Allen et al., 2009; O’Grady et al., 2012). Though the description provides an indication of carbon lost relative to that converted to biomass, the calculation also reflects the composition of the biomass. Seeds that make large amounts of lipid produce more CO_2_ as a part of fatty acid biosynthesis than seeds that predominantly make starch. Green seeds capitalize on photosynthesis to improve carbon efficiency (Schwender et al., 2004; Allen et al., 2009). Thus the CUE calculation must take into consideration the metabolic context that differs amongst seeds and tissues.

Analogously, the temporal dynamics of seed metabolism are self-evident from the dramatic change in seed appearance with development. Soybean seeds are green at R5 (Figure 2A) and are capable of productively using available sunlight (Ruuska et al., 2004; Borisjuk et al., 2005; Rolletschek et al., 2005; reviewed in Angelovici et al., 2010) due in part to the contribution of Rubisco-based CO_2_ fixation (Schwender et al., 2004; Allen et al., 2009). CUEs for stages in metabolism were calculated based on differences in composition and CO_2_ production over developmental stages, resulting in values of 90% and 76% between R5-R6 and R6-R7, respectively (Supplementary Table S3). Prior reports (Allen et al., 2009) that focused exclusively on seed filling indicated reasonable agreement with these early stages. During this time, the spike in CO_2_ production occurs when lipid production is high and seeds are starting to transition from green to yellow in color (Figure 2C). TCA cycle metabolism and elevated oxidative PPP were also supported during this interval based on related metabolite clustering analysis (Figure 5). Later in development, differences in CO_2_ production correlated with parallel drops in TCA cycle and OPPP metabolite levels (cluster 3 of Figure 4).

As seeds continue to develop, the CUE calculation is no longer applicable because the supply of exudate from the maternal plant is exhausted. Instead the balance of one reserve turned over should equate with production of other reserves and CO_2_, to avoid violation of mass conservation. Developmental staging showed that starch levels drop (Figure 2) to supply other needs including RFO and lipid biosynthesis. From R7 to maturation, the balance of carbon turned over as lipid and protein must account for new production of carbohydrates and CO_2_ generated. The reported changes in storage reserves (Supplementary Table 3) indicated a balance that was 94% closed. The production of some CO_2_ late in development may suggest a slight decrease in final seed biomass with desiccation; however, the change in seed weight was not statistically significant.

Interestingly, pertaining to RFO production, our results suggested biosynthesis at multiple locations, with carbon supplied from turned over storage reserves in the cotyledon and also as a result of the withering of the seed coat during dessication. ^13^C_12_ sucrose incubation (i.e., a precursor to raffinose) with seed coats indicated production of RFOs (Figure 7) and suggests that the reduction in seed coat biomass during maturation may be analogous to a senescence process where the carbohydrate in the seed coat is converted to RFOs at the surface of the cotyledon.

### Gluconeogenic activity is temporally synchronized with lipid degradation to supply turned over carbon for carbohydrate biosynthesis

Steady-state metabolic flux analysis using developing soybean seeds previously indicated that gluconeogenic metabolism does not occur when seeds receive adequate supplies of sugars (Allen et al., 2009). However, the composition of seed exudate late in development indicates that seeds do not receive an extensive supply of sugars from the maternal plant at these stages (Figure 4). As shown with ^13^C_3_ glycerol labeling experiments (Figure 5A), enzymes involved in shuttling carbon to hexose phosphates can operate effectively late in development so that carbon can be used to produce carbohydrates (RFOs and CWPs). ^13^C enrichment occured in intermediates of glycolysis (DHAP, PEP), hexose phosphates (G6P, G1P), and the nucleotide sugar UDPG, which are precursors to carbohydrate production (Figure 6). From the balance of biomass components (Supplementary Table 3), the CO_2_ loss of 3.3 mg (calculated) could come from PEPCK activity (61%; see Supplementary Table 3 for details) which was highest at R7.

The carbon required for PEPCK activity is likely obtained via the glyoxylate cycle, which utilizes acetyl-CoA derived from repeated deacylation of lipids beginning at R7 and continuing throughout the maturation phase by β-oxidation. The key enzymes required for glyoxylate cycle and β-oxidation, isocitrate lyase (ICL), malate synthase (MS), 3-ketoacyl-CoA thiolase (KAT), and the multifunctional enzyme (MFP) of β-oxidation, were previously shown to increase in activity during the maturation phase of embryo development in *Brassica napus*, characterized by lipid degradation (Chia et al., 2005). We observed that the glyoxylate levels increased over the course of development, peaking at R7 and R7.5 (cluster 4 of Figure 5, Supplementary Table 2). Further, activity of the glyoxylate cycle and gluconeogenesis are supported by prior measurements of transcript abundance over soybean development (Collakova et al., 2013; Li et al., 2015). The differences in carbon movement between the R6 stage of seed filling and inititation of maturation at R7 are stark and are summarized in Figure 8. Carbon received as sucrose via maternal contribution in R6 is channeled into lipid biosynthesis at R6. As the maternal contribution decreases between R6 to R7, starch turnover may contribute carbon to both lipid and oligosaccharide biosynthesis while TCA cycle operation sustains the energy required for oligosaccharide production. As the seeds reach the R7 maturation phase, turnover of lipids is initiated and the carbon from the glycerol backbones of lipids as well as degraded acyl chains is channeled into cell wall polysaccharides via gluconeogenic and β-oxidative pathways, respectively.

**Figure 8:**
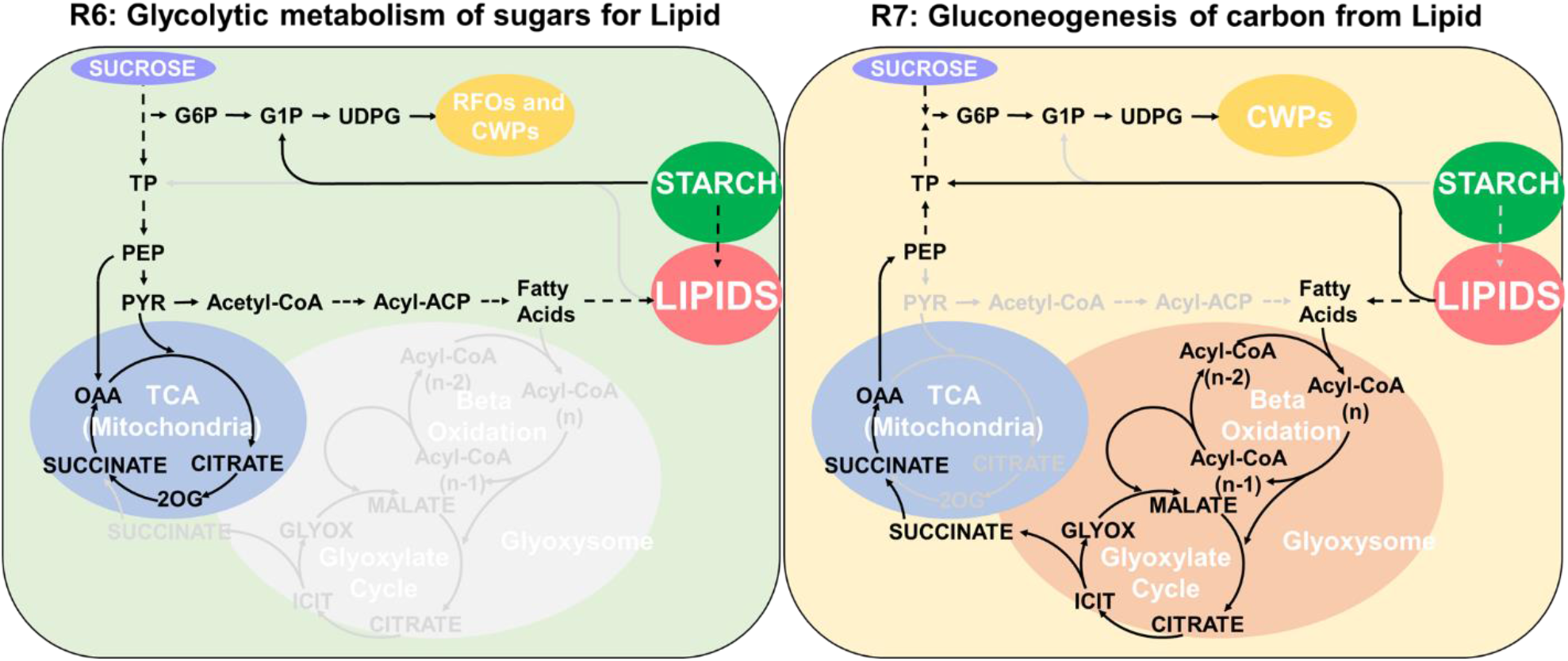
Proposed model for metabolic switch between R6 and R7 to shuttle carbon from starch and lipid breakdown to oligosaccharide and cell wall polysaccharide biosynthesis. As described in text, sources of carbon in R6 used for reserve production and energy metabolism are not present late in development and result in some storage reserves being turned over to support biosynthesis of others. Abbreviations: G6P, glucose 6-phosphate; G1P, glucose 1-phosphate; UDPG, uridine diphosphate glucose; TP, triose phosphate; PEP, phophoenol pyruvate; PYR, pyruvate; ACP, acyl carrier protein; OAA, oxaloacetic acid; 2OG, 2-oxoglutarate; RFO, raffinose family oligosaccharides; CWP, cell wall polysaccharides; TCA, tricarboxylic acid; GLYOX, glyoxylate; ICIT, isocitrate.

Recent efforts to improve seed quality have targeted lipases to reduce lipid breakdown late in development (Kanai et al., 2019) and RFO biosynthetic steps (Valentine et al., 2017; Hagely et al., 2020) to improve seed compositional traits without significant phenotypic consequences to maturation or germination. Carbohydrates as a whole (RFOs and CWPs) constitue ~40% of final seed composition and are an important sink and potentially rob carbon from the production of oil and protein. Hence, future engineering efforts for increased oil and protein, if focused on manipulating key carbon partitioning pathway nodes that consider all three biomass components, can be fruitful in breaking the perceived inverse correlation. Improved channeling of carbon from malate towards lipid using malic enzyme (Allen and Young, 2013) temporally and increasing the sink strength of developing seeds (Rolletschek et al., 2020) by manipulating the hormone status (Quoc Thien et al., 2016; Kambhampati et al., 2017) represent unrealized potential targets for future soybean improvement.

## Materials and Methods

### Plant growth conditions and tissue collection for *in planta* and seed coat exudate measurements

Soybean cultivar, Williams 82, was grown under greenhouse conditions as previously described (Kambhampati et al., 2019). Germinating seeds were transferred to one-gallon pots containing Fafard 4M and grown at 25°C-27°C/21°C-23°C day/night temperatures with greater than 35% humidity and sunlight supplemented by approximately 400-1000 Wm^−2^ to establish a 14 hr day, 10 hr night photoperiod. Plants were watered daily and received Jack’s 15-16-17 (JR Peters) fertilizer three times a week. Developing seeds were grouped based on fresh weight and visual appearance to determine the developmental stage (Figure 1). At the time of harvest, seeds that were used as controls representing *in planta* conditions were dissected from pods, the seed coat was removed and cotyledons were flash frozen with liquid nitrogen and stored at −80°C until further use. Cotyledons collected from a single plant were treated as a single biological replicate for all stages of development. Tissue that belonged to each replicate was ground individually using machined home-made stainless-steel hammer-crushing pestle and mortar design. Ground and frozen tissue was then lyophilized and aliquoted for individual biomass component and metabolite measurements.

For exudate experiments an isotonic solution of 20 mM ammonium acetate pH 6.8 was placed on the interior side of excised seed coats and pipeted up and down repeatedly for 10 seconds. In addition, the surface of the corresponding cotyledon was rinsed with the same solution to collect any surface contents. The extracts from each stage were dried using a speed vacuum centrifuge and resuspeneded in water and filtered using 0.45 μm cellulose acetate centrifuge filters (costar®, Corning Inc.) prior to metabolite measurements using LC-MS/MS.

### Moisture content, fresh weight, dry weight, and net CO_2_ measurements

Moisture content was determined using fresh weight and dry weight measurements. Seeds were weighted immediately after harvesting to obtain fresh weights, sliced and dried in an oven. Dried seeds were measured at least three times over the course of several weeks to ensure no moisture remained and the weights plateauted. CO_2_ measurements were taken from whole soybean cotyledons, after excising the pod walls and seed coats, using a LI-COR 6400 with an attached insect respiration chamber (#6400-89) following manufacturers protocol. Five replicates with three cotyledons each and ten measurements were used with readings taken every 20 to 30 seconds. 30 μE of light was maintained throughout the measurement period to simulate light received by the cotyledons within the pods (Allen et al., 2009).

### Normal seed RFO soybean cultivar ‘Jack’ and the ultra-low RFO derivative ‘Jack *rs2 rs3*’

An ultra-low RFO version of soybean cultivar ‘Jack’ (Jack *rs2 rs3*) was developed by backcrossing variant alleles of the raffinose synthase 2 (*rs2*) and raffinose synthase 3 (*rs3*) genes into Jack (Hagely et al., 2020). The ultra-low RFO phenotype has been defined as raffinose and stachyose content less than 0.70% of seed dry weight (Hagely et al., 2013; Schillinger JA, 2013, 2018).

### ^13^C labeled culture system set up and conditions

For culturing system development, cotyledons from specific stages were excised from seed coats under sterile conditions and immediately placed flat face down for each cotyledonary half into 300 μL of sterile culture medium in a 24-well plate. A modified Linsmaier and Skoog medium (Thompson et al., 1977; Hsu and Obendorf, 1982) with Gamborg’s vitamins (Sigma) and 5 mM MES buffer adjusted to pH 5.8 was used as the culture medium and contained 200 mM U-^13^C_6_ glucose as the exclusive carbon source. Culturing was performed under 30 – 35 μE continuous light at 26°C, consistent with Allen et al. (2009) and tissue was collected at 5, 10, and 30 mins, in triplicates, for all time course studies described. Untreated samples were also taken in triplicates for 0 timepoint (*in-planta*) measurements. At the conclusion of labeling experiments, the metabolism was quenched by a very brief rinse of the cotyledon surface with water prior to slicing off layers of the cotyledon to assess label uptake, metabolism, and heterogeneity (see Supplementary Data 1 for details). Slices were rapidly frozen in liquid nitrogen and stored at −80°C until extraction.

To investigate carbon turnover from lipids to carbohydrates, we substituted U-^13^C_6_ glucose with ^13^C_3_ glycerol (15 mM) as the sole source of carbon. ^13^C_12_ Sucrose (100 mM) was used as the carbon source for data presented in Figure 7. The salts and vitamins in all individual labelling experiments remained the same as above. The bottom slice of labeled seeds was used for metabolite extraction and measurements of isotopologue distribution (see Supplementary Data 1 for rationale).

### RNA extraction and transcript analysis for verification of culturing system

Total RNA was extracted from soybean seed slices using the RNeasy Plant Mini Kit (Qiagen) according to the supplier’s instructions. cDNA was synthesized using the qScript cDNA SuperMix (Quantabio) from 1 μg total RNA previously treated with DNase I (Merck). For droplet generation, 20 μl of PCR reaction (cDNA, primers and the Bio-Rad ddPCR supermix) and 70 μl of droplet generation oil were transferred to the middle and to the bottom rows respectively of a DG8™ Cartridge before insertion into a QX200 Droplet Generator. The genes used and the primer sequences are included in Supplementary Table S4. Droplets were transferred to a 96-well plate for PCR amplification in a thermal cycler C1000 Touch™. The cycling protocol was 95 °C enzyme activation for 5 min followed by 40 cycles of a two-step cycling protocol of 95 C° for 30 seconds and Tm (Supplementary Table S4) for 1 min, then 4 °C for 5 min and 90 °C for 5min. Following PCR amplification, the plate containing the droplets was placed in a QX200 droplet reader. Droplet digital PCR (ddPCR) data was analysed with Bio-Rad QuantaSoft Analysis Pro Software. The *Glycine max* ATP synthase subunit 1 (Glyma12g02310) and SKIP16 (Glyma12g05510) were used as internal references (Hu et al., 2009).

### Extraction of polar and non-polar metabolites

Metabolite extraction was carried out following the protocol described in Czajka et al. (2020)) and Kambhampati et al. (2019)) with a few modifications. Briefly, the stored samples were removed from −80°C and two metal beads were added to each tube along with 1 ml 7:3 methanol/chloroform (−20°C) and a PIPES (piperazine-N,N’-bis[2-ethanesulfonic acid]), norvaline, and ribitol mixed standard. Samples were kept on ice throughout extraction unless otherwise noted. Samples were pulverized using a ball mill at 28 Hz for 5 minutes or until fully ground. The mixtures were then incubated at −20°C for 3 hrs, with intermittent vortexing to ensure complete extraction. 500 μL of ddH_2_O (4°C) was added to each sample and vortexed vigorously before being centrifuged at 14,000 rpm at 4°C for 10 minutes during which the samples phase-separated. The upper aqueous phase containing water-soluble metabolites was transferred to a 1.5 mL eppindorf tube with a 0.45 μm centrifugal filter (Costar®, Corning Inc.) and spun at 14,000 rpm at 4°C for 2 mins. This solution was then transferred to glass vials (Agilent, Xpertek) for LC-MS/MS analysis to detect soluble sugars, free amino acids and sugar phosphates.

### Quantification of proteins, lipids, and starch

3-5 mg of ground lyophilized tissue was subjected to liquid hydrolysis and protein was measured using amino acid compositional analysis as described in Kambhampati et al. (2019). In brief, 20 μL of 1 mM cell free 13C-labelled amino acid standard mix (Sigma) was added to the protein pellet and dried using a speed vacuum centrifuge. 50 μL of 4M methanesulphonic acid containing 0.2% tryptamine was added to this dried pellet and incubated at 110°C for 22 hours. Upon completion of hydrolysis, the samples were neutralized using 50 μL of 4M sodium hydroxide, briefly vortexed and dried. Upon drying, the samples were resuspended in 1 mL ultra pure water and vortexed to recover the hydrolyzed amino acids and then filtered using 0.45 μM centrifugal filters. Amino acids were detected using LC-MS/MS (described below) and quantified using isotopic dilution based on peak areas obtained from known concentrations of internal standards. The sum of the concentrations of all 20 amino acids, in milligrams, was used to establish the concentration of protein (Supplementary Table S5).

Analysis of lipid content was carried out according to an adapted version of the method described in Allen and Young (2013)) by converting total lipids into Fatty Acid Methyl Esters (FAMEs). In brief, freshly prepared 5% sulfuric acid:methanol (v/v) was added to ~20 mg of ground lyophilized tissue along with 25 ul 0.2% butylated hydroxytoluene (BHT) in methanol to prevent oxidation and two internal standards, triheptadecanoin and tripentadecanoin, before heating at 110°C for 3 hours, vortexing hourly. After cooling to room temperature, 0.9% NaCl (w/v) was added to each sample to quench the reaction. The FAMEs were then extracted using hexane and quantified by gas chromatography-flame ionization detection (GC-FID) using a DB23 column (30 m, 0.25-mm i.d., 0.25-μm film; J&W Scientific). The GC was operated in a split mode (30:1). The flame ionization detector was operated with a temperature of 250°C with an oven temperature ramp profile from 180°C to 260°C at a rate of 20°C min^−1^ followed by a hold time of 7 mins. Comparisons of peak areas to the two internal standards were used for quantification.

Starch measurements were performed on ~20 mg of ground lyophilized tissue, directly without prior extraction. Total starch content in cotyledons over reproductive development was determined, in triplicates, using the Megazyme starch assay kit (Megazyme International Ireland), using the AOAC Official Method 996.11 (Approved Methods of the AACC,(McCleary et al., 1997; McCleary et al., 2019)) modified to adjust the final assay volume for 96-well plate reader compatibility. Briefly, the ground lyophilized tissue was washed twice with 80% ethanol at 85°C prior to heating at 110°C for 10 min with DMSO. The samples were then treated with α-amylase at 110°C for 12 minutes (vortexing every 4 min) followed by amyloglucosidase at 50°C for 1 hour. The samples were then centrifuged, supernatant collected, and incubated with the GOPOD reagent at 50°C for 20 min. The absorbace at 510 nm was measured using a spectrophotometer, and the starch content was determined by comparison with a standard curve generated using a serial dilution of starch standards treated the same way as biological samples. Quantities of all biomass component measurements presented in Figure 2 are provided as Supplementary Table S6.

### Quantification of soluble sugars, amino acids, and sugar phosphates using HPLC-MS/MS

Sugars and sugar phosphates were analyzed from the water-soluble fraction using a Shimadzu (UFLCXR) HPLC system connected to an AB Sciex triple quadrupole MS equipped with Turbo V™ electrospray ionization (ESI) source using the method described in Czajka et al. (2020)). Negative ion mode was used to monitor sugar and sugar phosphate fragments. A 5 μl sample was injected on the Infinity Lab poroshell 120 Z-HILIC column (2.7 μm, 100 x 2.1 mm; Agilent technologies, Santa Clara, CA, USA) and the metabolites were eluted with an increasing gradient of acetonitrile: 10 mM ammonium acetate (90:10 v/v) and 5 μm medronic acid, pH 9.0 (A) and 10 mM ammonium acetate in water, pH 9.0 (B). The flow rate was 0.25 mL/min. Sugars and sugar phosphates were separated using a binary gradient of 95-70% B over 8 minutes then to 50% B over the next 4 min followed by a hold at 25% B for 1.5 min. The gradient was then decreased to 30% B over 0.5 min followed by a hold for 1 min before returning to 95% B to re-equilibrating the column for 6 min. The HPLC eluent was introduced into an electrospray ionization source with the following conditions: ion spray voltage, 4.5 kV (ESI-); ion source temperature, 550°C; source gas 1, 45 psi; source gas 2, 40 psi; curtain gas, 35 psi and entrance potential, 10. Ions were detected and monitored using a targeted MRM approach with the parameters optimized by direct infusions, provided in supplementary Table S7, for accurate quantification. The value for entrance potential was default (−10) for all analytes. For absolute quantification, data were analyzed using the quantitation wizard available in Analyst (v. 1.6.2) software (AB SCIEX, Concord, Canada). Metabolite concentrations were calculated based on a calibration curves. Recoveries were assessed using ribitol and PIPES as internal standards for sugars and sugar phosphates, respectively.

Amino acids were measured using the same instrumentation and column as described above, with the following changes in mobile phases, gradient and ionization conditions; Mobile phase A consisted of 20mM ammonium formate in water, pH 3.0, and B was composed of 90% acetonitrile and 10% water with a final concentration of 20 mM ammonium formate, pH 3. 3 μL from each sample were injected and a flow rate of 0.25 μL was used for separation of amino acids on the HPLC column. A binary gradient composed of 100-90% B over 2 minutes, 90-50% B over the next 6 minutes followed by returning to 100% B over 30 seconds and re-equilibration of 5.5 minutes was used to separate the analytes. The HPLC eluent was introduced into an electrospray ionization source with the following conditions: ion spray voltage, 4.5 kV (ESI+ and ESI-); ion source temperature, 400°C; source gas 1, 45; source gas 2, 40; curtain gas, 35 and entrance potential, 10. Ions were detected and monitored using a targeted MRM approach using parameters included within Supplementary Table S7. Data were analyzed similar to sugars and sugar phosphates described above, except novaline was used as an internal standard to assess recoveries. All statistical analysis and data visualization were performed using Microsoft Excel (2013) or R programming language (R CoreTeam, 2013) using base functions and the package ggplot2 (Wickham, 2016).

### Quantification of Isotopologue abundance and average labeling

The LC-MS/MS conditions used for label detection are identical to the ones described above with the exception of MRM transitions used (Supplementary Table S8) that were selected based on Kappelmann et al. (2017). Peaks were manually integrated and the natural abundance was corrected using the R package IsoCorrectoR (Heinrich et al., 2018). Fractional enrichment of the corrected isotopologues (M0-Mn) obtained from IsoCorrectoR was used to calculate average labeling. Average labeling was calculated as described in Buescher et al. (2015) using the following equation;

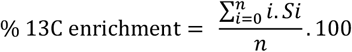

where *i* denotes the mass isotopologue, *n* is the number of possible ^13^C carbons, and *S* is the fraction of the labeled isotopologue. For detailed calculations see Supplementary Table S9.

### PEPCK enzyme activity assay

PEPCK enzyme activity was measured using the method of Walker et al. (1999) detailed on protocols.org (Osorio et al., 2014). Briefly, crude protein was extracted from 100-250mg FW of seed tissue in a buffer containing 0.5M bicine-KOH (pH 9.0), 0.2M KCl, 3mM EDTA, 5% (w/v) PEG-4000, 25mM DTT, and 0.4% bovine serum albumin. The extract was centrifuged for 20 minutes at 13,000 x g at 4°C, and the supernatant was added to a buffer containing 0.5M bicine-KOH (pH 9.0), 3mM EDTA, 55% (w/v) PEG-4000, and 25mM DTT, then incubated for 10 minutes on ice and centrifuged at 13,000 x g at 4°C for 20 minutes. The supernatant was discarded, and the pellet was resuspended in 10mM bicine-KOH (pH 9.0) containing 25mM DTT. PEPCK activity was measured in the direction of the carboxylation reaction by coupling the reaction with malate dehydrogenase (EC 1.1.1.37; Sigma Aldrich 10127914001) and following the oxidation of NADH at 340nm at room temperature using a spectrophotometer (SpectraMax M2^e^, Molecular Devices). Total protein was measured using the protein extract for PEPCK activity using Bradford reagent (Millipore Sigma; Cat: B6916) and a standard curve using commercial bovine serum albumin standards (Thermo Fisher, cat: 23208).

## Supplementary material

**Supplementary data 1:** Establishing and assessing short time pulse labeling conditions for non-perturbed *in planta* temporal assessment of seed development

**Supplementary Figure S1:** ^13^C_6_ glucose labeling using cotyledons of R7 seeds for establishing the culturing system.

**Supplementary Table S1:** Composition of seed coat exudate, quantities represented in nmol per seed

**Supplementary Table S2:** Total pool size (accurate quantities) of the metabolites detected via LC-MS/MS

**Supplementary Table S3:** Temporal changes in carbon conversion efficiency and loss of CO_2_ as PEPCK

**Supplementary Table S4:** Primers used for droplet digital PCR and their annealing temperatures

**Supplementary Table S5:** Amino acid concentration (mg seed^−1^) from hydrolyzed protein at different developmental stages

**Supplementary Table S6:** Biomass component measurements for soybean seed developmental stages

**Supplementary Table S7:** MRM parameters used for absolute quantification

**Supplementary Table S8:** MRM parameters used for isotope labeling experiments

**Supplementary Table S9:** Calculations for isotopologue distribution and average labeling

## Acknowledgements

The authors would like to thank the director of proteomics and mass spectrometry (PMSF) facility at the Donald Danforth Plant Science Center (DDPSC), Dr. Bradley S. Evans for his support and advice throughout this work. In addition, the authors acknowledge the Plant Growth Facility (PGF) at DDPSC for their help with plant growth and maintenance. The authors declare no conflicts of interest.

## References

Adams CA, Fjerstad MC, Rinne RW (1983) Characteristics of Soybean Seed Maturation: Necessity for Slow Dehydration1. Crop Science 23: 265–267

Allen DK (2016) Quantifying plant phenotypes with isotopic labeling & metabolic flux analysis. Current Opinion in Biotechnology 37: 45–52

Allen DK, Bates PD, Tjellström H (2015) Tracking the metabolic pulse of plant lipid production with isotopic labeling and flux analyses: Past, present and future. Progress in Lipid Research 58: 97–120

Allen DK, Ohlrogge JB, Shachar-Hill Y (2009) The role of light in soybean seed filling metabolism. The Plant Journal 58: 220–234

Allen DK, Young JD (2013) Carbon and Nitrogen Provisions Alter the Metabolic Flux in Developing Soybean Embryos. Plant Physiology 161: 1458–1475

Alonso AP, Goffman FD, Ohlrogge JB, Shachar-Hill Y (2007) Carbon conversion efficiency and central metabolic fluxes in developing sunflower (Helianthus annuus L.) embryos. The Plant Journal 52: 296–308

Angelovici R, Galili G, Fernie AR, Fait A (2010) Seed desiccation: a bridge between maturation and germination. Trends in Plant Science 15: 211–218

Assefa Y, Bajjalieh N, Archontoulis S, Casteel S, Davidson D, Kovács P, Naeve S, Ciampitti IA (2018) Spatial Characterization of Soybean Yield and Quality (Amino Acids, Oil, and Protein) for United States. Scientific Reports 8: 14653

Bates P, Browse J (2012) The Significance of Different Diacylgycerol Synthesis Pathways on Plant Oil Composition and Bioengineering. Frontiers in Plant Science 3

Baud S, Boutin J-P, Miquel M, Lepiniec L, Rochat C (2002) An integrated overview of seed development in Arabidopsis thaliana ecotype WS. Plant Physiology and Biochemistry 40: 151–160

Baud S, Graham IA (2006) A spatiotemporal analysis of enzymatic activities associated with carbon metabolism in wild-type and mutant embryos of Arabidopsis using in situ histochemistry. The Plant Journal 46: 155–169

Baud S, Lepiniec L (2009) Regulation of de novo fatty acid synthesis in maturing oilseeds of Arabidopsis. Plant Physiology and Biochemistry 47: 448–455

Baud S, Wuillème S, To A, Rochat C, Lepiniec L (2009) Role of WRINKLED1 in the transcriptional regulation of glycolytic and fatty acid biosynthetic genes in Arabidopsis. The Plant Journal 60: 933–947

Borisjuk L, Nguyen TH, Neuberger T, Rutten T, Tschiersch H, Claus B, Feussner I, Webb AG, Jakob P, Weber H, Wobus U, Rolletschek H (2005) Gradients of lipid storage, photosynthesis and plastid differentiation in developing soybean seeds. New Phytologist 167: 761–776

Buescher JM, Antoniewicz MR, Boros LG, Burgess SC, Brunengraber H, Clish CB, DeBerardinis RJ, Feron O, Frezza C, Ghesquiere B, Gottlieb E, Hiller K, Jones RG, Kamphorst JJ, Kibbey RG, Kimmelman AC, Locasale JW, Lunt SY, Maddocks ODK, Malloy C, Metallo CM, Meuillet EJ, Munger J, Nöh K, Rabinowitz JD, Ralser M, Sauer U, Stephanopoulos G, St-Pierre J, Tennant DA, Wittmann C, Vander Heiden MG, Vazquez A, Vousden K, Young JD, Zamboni N, Fendt S-M (2015) A roadmap for interpreting 13C metabolite labeling patterns from cells. Current Opinion in Biotechnology 34: 189–201

Chapman KD, Dyer JM, Mullen RT (2012) Biogenesis and functions of lipid droplets in plants: Thematic Review Series: Lipid Droplet Synthesis and Metabolism: from Yeast to Man. Journal of Lipid Research 53: 215–226

Chia TYP, Pike MJ, Rawsthorne S (2005) Storage oil breakdown during embryo development of Brassica napus (L.). Journal of Experimental Botany 56: 1285–1296

Clemente TE, Cahoon EB (2009) Soybean Oil: Genetic Approaches for Modification of Functionality and Total Content. Plant Physiology 151: 1030–1040

Collakova E, Aghamirzaie D, Fang Y, Klumas C, Tabataba F, Kakumanu A, Myers E, Heath LS, Grene R (2013) Metabolic and Transcriptional Reprogramming in Developing Soybean (Glycine max) Embryos. Metabolites 3: 347–372

Czajka JJ, Kambhampati S, Tang YJ, Wang Y, Allen DK (2020) Application of Stable Isotope Tracing to Elucidate Metabolic Dynamics During Yarrowia lipolytica α-Ionone Fermentation. iScience 23: 100854

Dierking EC, Bilyeu KD (2009) Raffinose and stachyose metabolism are not required for efficient soybean seed germination. Journal of Plant Physiology 166: 1329–1335

Dyer JM, Stymne S, Green AG, Carlsson AS (2008) High-value oils from plants. The Plant Journal 54: 640–655

Eastmond PJ, Germain V, Lange PR, Bryce JH, Smith SM, Graham IA (2000) Postgerminative growth and lipid catabolism in oilseeds lacking the glyoxylate cycle. Proceedings of the National Academy of Sciences 97: 5669–5674

Eastmond PJ, Graham IA (2001) Re-examining the role of the glyoxylate cycle in oilseeds. Trends in Plant Science 6: 72–78

Egli DB, Bruening WP (2001) Source-sink Relationships, Seed Sucrose Levels and Seed Growth Rates in Soybean. Annals of Botany 88: 235–242

Fabre F, Planchon C (2000) Nitrogen nutrition, yield and protein content in soybean. Plant Science 152: 51–58

Fait A, Angelovici R, Less H, Ohad I, Urbanczyk-Wochniak E, Fernie AR, Galili G (2006) Arabidopsis Seed Development and Germination Is Associated with Temporally Distinct Metabolic Switches. Plant Physiology 142: 839–854

Gawłowska M, Święcicki W, Lahuta L, Kaczmarek Z (2017) Raffinose family oligosaccharides in seeds of Pisum wild taxa, type lines for seed genes, domesticated and advanced breeding materials. Genetic Resources and Crop Evolution 64: 569–578

Gifford RM, John HT (1985) Sucrose Concentration at the Apoplastic Interface between Seed Coat and Cotyledons of Developing Soybean Seeds. Plant Physiology 77: 863–868

Gomes CI, Obendorf RL, Horbowicz M (2005) myo-Inositol, D-chiro-Inositol, and D-Pinitol Synthesis, Transport, and Galactoside Formation in Soybean Explants. Crop Science 45: 1312–1319

Hagely KB, Jo H, Kim J-H, Hudson KA, Bilyeu K (2020) Molecular-assisted breeding for improved carbohydrate profiles in soybean seed. Theoretical and Applied Genetics 133: 1189–1200

Hagely KB, Palmquist D, Bilyeu KD (2013) Classification of Distinct Seed Carbohydrate Profiles in Soybean. Journal of Agricultural and Food Chemistry 61: 1105–1111

Heinrich P, Kohler C, Ellmann L, Kuerner P, Spang R, Oefner PJ, Dettmer K (2018) Correcting for natural isotope abundance and tracer impurity in MS-, MS/MS-and high-resolution-multiple-tracer-data from stable isotope labeling experiments with IsoCorrectoR. Scientific Reports 8: 17910

Hernández-Sebastià C, Marsolais F, Saravitz C, Israel D, Dewey RE, Huber SC (2005) Free amino acid profiles suggest a possible role for asparagine in the control of storage-product accumulation in developing seeds of low- and high-protein soybean lines. Journal of Experimental Botany 56: 1951–1963

Hsu FC, Bennett AB, Spanswick RM (1984) Concentrations of Sucrose and Nitrogenous Compounds in the Apoplast of Developing Soybean Seed Coats and Embryos. Plant Physiology 75: 181–186

Hsu FC, Obendorf RL (1982) Compositional analysis of in vitro matured soybean seeds. Plant Science Letters 27: 129–135

Hu R, Fan C, Li H, Zhang Q, Fu Y-F (2009) Evaluation of putative reference genes for gene expression normalization in soybean by quantitative real-time RT-PCR. BMC Molecular Biology 10: 93

Kambhampati S, Aznar-Moreno JA, Hostetler C, Caso T, Bailey SR, Hubbard AH, Durrett TP, Allen DK (2019) On the Inverse Correlation of Protein and Oil: Examining the Effects of Altered Central Carbon Metabolism on Seed Composition Using Soybean Fast Neutron Mutants. Metabolites 10: 18

Kambhampati S, Kurepin LV, Kisiala AB, Bruce KE, Cober ER, Morrison MJ, Emery RJN (2017) Yield associated traits correlate with cytokinin profiles in developing pods and seeds of field-grown soybean cultivars. Field Crops Research 214: 175–184

Kambhampati S, Li J, Evans BS, Allen DK (2019) Accurate and efficient amino acid analysis for protein quantification using hydrophilic interaction chromatography coupled tandem mass spectrometry. Plant Methods 15: 46

Kanai M, Yamada T, Hayashi M, Mano S, Nishimura M (2019) Soybean (Glycine max L.) triacylglycerol lipase GmSDP1 regulates the quality and quantity of seed oil. Scientific Reports 9: 8924

Kappelmann J, Klein B, Geilenkirchen P, Noack S (2017) Comprehensive and accurate tracking of carbon origin of LC-tandem mass spectrometry collisional fragments for 13C-MFA. Analytical and Bioanalytical Chemistry 409: 2309–2326

Kosina SM, Castillo A, Schnebly SR, Obendorf RL (2009) Soybean seed coat cup unloading on plants with low-raffinose, low-stachyose seeds. Seed Science Research 19: 145–153

Kuo TM, VanMiddlesworth JF, Wolf WJ (1988) Content of raffinose oligosaccharides and sucrose in various plant seeds. Journal of Agricultural and Food Chemistry 36: 32–36

Leprince O, Pellizzaro A, Berriri S, Buitink J (2016) Late seed maturation: drying without dying. Journal of Experimental Botany 68: 827–841

Li L, Hur M, Lee J-Y, Zhou W, Song Z, Ransom N, Demirkale CY, Nettleton D, Westgate M, Arendsee Z, Iyer V, Shanks J, Nikolau B, Wurtele ES (2015) A systems biology approach toward understanding seed composition in soybean. BMC Genomics 16: S9

Licht M (2014) Soybean Growth and Development. In, Vol 2019, Iowa State University Extension and Outreach

Lin W, Oliver DJ (2008) Role of triacylglycerols in leaves. Plant Science 175: 233–237

McCleary BV, Charmier LMJ, McKie VA (2019) Measurement of Starch: Critical Evaluation of Current Methodology. Starch – Stärke 71: 1800146

McCleary BV, Gibson TS, Mugford DC, Collaborators (1997) Measurement of Total Starch in Cereal Products by Amyloglucosidase-α-Amylase Method: Collaborative Study. Journal of AOAC INTERNATIONAL 80: 571–579

Mello Filho OLd, Sediyama CS, Moreira MA, Reis MS, Massoni GA, Piovesan ND (2004) Grain yield and seed quality of soybean selected for high protein content. Pesquisa Agropecuária Brasileira 39: 445–450

Naeve SL (2005) Soybean growth stages. In, Vol 2019, University of Minnesota Extension

O’Grady J, Schwender J, Shachar-Hill Y, Morgan JA (2012) Metabolic cartography: experimental quantification of metabolic fluxes from isotopic labelling studies. Journal of Experimental Botany 63: 2293–2308

Osorio S, Vallarino JG, Szecowka M, Ufaz S, Tzin V, Angelovici R, Galili G, Aarabi F (2014) Extraction and Measurement the Activities of Cytosolic Phosphoenolpyruvate Carboxykinase (PEPCK) and Plastidic NADP-dependent Malic Enzyme (ME) on Tomato (Solanum lycopersicum). Bio-protocol 4: e1122

Patil G, Mian R, Vuong T, Pantalone V, Song Q, Chen P, Shannon GJ, Carter TC, Nguyen HT (2017) Molecular mapping and genomics of soybean seed protein: a review and perspective for the future. Theoretical and Applied Genetics 130: 1975–1991

Pipolo AE, Sinclair TR, Camara GMS (2004) Protein and oil concentration of soybean seed cultured in vitro using nutrient solutions of differing glutamine concentration. Annals of Applied Biology 144: 223–227

Quoc Thien N, Anna K, Peter A, Emery RJN, Suresh N (2016) Soybean Seed Development: Fatty Acid and Phytohormone Metabolism and Their Interactions. Current Genomics 17: 241–260

Rainbird RM, Thorne JH, Hardy RWF (1984) Role of Amides, Amino Acids, and Ureides in the Nutrition of Developing Soybean Seeds. Plant Physiology 74: 329–334

Raymond R, Spiteri A, Dieuaide M, Gerhardt B, Pradet A (1992) Peroxisomal beta – oxidation of fatty acids and citrate formation by a particulate fraction from early germinating sunflower seeds. 30: 153–161

Rolletschek H, Radchuk R, Klukas C, Schreiber F, Wobus U, Borisjuk L (2005) Evidence of a key role for photosynthetic oxygen release in oil storage in developing soybean seeds. New Phytologist 167: 777–786

Rolletschek H, Schwender J, Konig C, Chapman KD, Romsdahl T, Lorenz C, Braun HP, Denolf P, Van Audenhove K, Munz E, Heinzel N, Ortleb S, Rutten T, McCorkle S, Borysyuk T, Guendel A, Shi H, Vander Auwermeulen M, Bourot S, Borisjuk L (2020) Cellular Plasticity in Response to Suppression of Storage Proteins in the Brassica napus Embryo. Plant Cell 32: 2383–2401

Rolletschek H, Weber H, Borisjuk L (2003) Energy Status and Its Control on Embryogenesis of Legumes. Embryo Photosynthesis Contributes to Oxygen Supply and Is Coupled to Biosynthetic Fluxes. Plant Physiology 132: 1196–1206

Ruuska SA, Schwender J, Ohlrogge JB (2004) The Capacity of Green Oilseeds to Utilize Photosynthesis to Drive Biosynthetic Processes. Plant Physiology 136: 2700–2709

Salon C, Raymond P, Pradet A (1988) Quantification of carbon fluxes through the tricarboxylic acid cycle in early germinating lettuce embryos. Journal of Biological Chemistry 263: 12278–12287

Sánchez-Mata MC, Peñuela-Teruel MJ, Cámara-Hurtado M, Díez-Marqués C, Torija-Isasa ME (1998) Determination of Mono-, Di-, and Oligosaccharides in Legumes by High-Performance Liquid Chromatography Using an Amino-Bonded Silica Column. Journal of Agricultural and Food Chemistry 46: 3648–3652

Schillinger JA DE, Bilyeu KD (2013) Soybeans having high germination rates and ultra-low raffinose and stachyose content. In USPTO, ed, Vol 8471107, USA

Schillinger JA DE, Bilyeu KD (2018) Soybeans having high germination rates and ultra-low raffinose and stachyose content. In USPTO, ed, Vol 10081814, USA

Schwender J, Goffman F, Ohlrogge JB, Shachar-Hill Y (2004) Rubisco without the Calvin cycle improves the carbon efficiency of developing green seeds. Nature 432: 779–782

Schwender J, Ohlrogge JB (2002) Probing in Vivo Metabolism by Stable Isotope Labeling of Storage Lipids and Proteins in Developing (Brassica napus) Embryos. Plant Physiology 130: 347–361

Schwender J, Shachar-Hill Y, Ohlrogge JB (2006) Mitochondrial Metabolism in Developing Embryos of Brassica napus. Journal of Biological Chemistry 281: 34040–34047

Singh SK, Barnaby JY, Reddy VR, Sicher RC (2016) Varying Response of the Concentration and Yield of Soybean Seed Mineral Elements, Carbohydrates, Organic Acids, Amino Acids, Protein, and Oil to Phosphorus Starvation and CO2 Enrichment. Frontiers in Plant Science 7

Team RC (2013) A language and environment for statistical computing. In R Foundation for Statistical Computing, Vienna, Austria

Thompson JF, Madison JT, Muenster A-ME (1977) In vitro Culture of Immature Cotyledons of Soya Bean (Glycine max L. Merr.). Annals of Botany 41: 29–39

Truong Q, Koch K, Yoon JM, Everard JD, Shanks JV (2013) Influence of carbon to nitrogen ratios on soybean somatic embryo (cv. Jack) growth and composition. Journal of Experimental Botany 64: 2985–2995

Tschiersch H, Borisjuk L, Rutten T, Rolletschek H (2011) Gradients of seed photosynthesis and its role for oxygen balancing. Biosystems 103: 302–308

Valentine MF, De Tar JR, Mookkan M, Firman JD, Zhang ZJ (2017) Silencing of Soybean Raffinose Synthase Gene Reduced Raffinose Family Oligosaccharides and Increased True Metabolizable Energy of Poultry Feed. Frontiers in Plant Science 8

Walker RP, Chen Z-H, Técsi LI, Famiani F, Lea PJ, Leegood RC (1999) Phosphoenolpyruvate carboxykinase plays a role in interactions of carbon and nitrogen metabolism during grape seed development. Planta 210: 9–18

Wickham H (2016) ggplot2: Elegant Graphics for Data Analysis. Springer Publishing Company, Incorporated

